# Autism-associated mutation in Hevin/Sparcl1 induces endoplasmic reticulum stress through structural instability

**DOI:** 10.1101/2022.04.12.488096

**Authors:** Takumi Taketomi, Takunori Yasuda, Rikuri Morita, Jaehyun Kim, Yasuteru Shigeta, Cagla Eroglu, Ryuhei Harada, Fuminori Tsuruta

## Abstract

Hevin is a secreted extracellular matrix protein that is encoded by *SPARCL1* gene. Recent studies show that Hevin plays an important role in regulating synaptogenesis and synaptic plasticity. Mutations in *SPARCL1* gene increase the risk of autism spectrum disorder (ASD). However, the molecular basis of how mutations in *SPARCL1* increase the risk of ASD has not been fully understood. In this study, we show that one of *SPARCL1* mutations associated with ASD impairs normal Hevin secretion. We identified Hevin mutants lacking the EF-hand motif through analyzing ASD-related mice with vulnerable spliceosome functions. Hevin deletion mutants accumulate in the ER, leading to the activation of unfolded protein responses. We also found that a single amino acid substitution of Trp^647^ with Arg in the EF-hand motif associated with a familial case of ASD causes a similar phenotype with the EF-hand deletion mutant. Importantly, molecular dynamics (MD) simulation revealed that this single amino acid substitution triggers exposure of hydrophobic amino acid to the surface, increasing the binding of Hevin with a molecular chaperon, BIP. Taken together, these data suggest that the integrity of EF-hand motif in Hevin is crucial for proper folding and ASD-related mutation impairs an export of Hevin from the endoplasmic reticulum (ER). Our data provide a novel mechanism linking a point mutation in *SPARCL1* gene to the molecular and cellular characteristics involved in ASD.

## Introduction

Autism spectrum disorder (ASD) is a prevalent neurodevelopmental disorder characterized by impaired social interaction and behaviors. The congenital genetic defect is one of the causes of ASD. So far, many genes associated with ASD have been identified by several genome-wide association studies^1-5^. Mostly, these mutations are found in the genes encoding synaptic proteins, such as ion channels, neurotransmitters, transporters, scaffold proteins, neurotrophic factors, and cell adhesion molecules in the pre- and post-synapses. Hence, perturbed synaptic connectivity caused by mutations is highly linked to the pathogenesis of ASD.

Several studies have reported that the amino acid substitution of synaptic proteins accumulates in the endoplasmic reticulum (ER) and causes abnormal trafficking to the proper sites, leading to an increase in the seriousness of ASD. These substitutions impair protein folding and frequently activate unfolded protein response (UPR). For instance, the ASD-associated mutant of Neuroligin (NL) 4 in which Arg^87^ is replaced with Trp accumulates in the ER and does not transport to the synaptic site^6^. The ASD-associated γ-aminobutyric acid transporter 1 (GAT-1) mutant, in which Pro^361^ is replaced with Thr, is prone to localize in the ER^7^. Moreover, missense mutations of novel ASD-associated transmembrane proteins, a cell adhesion molecule-1 (CADM1), and contactin-associated protein-like 2 (CASPR2) are also localized in the ER and upregulates ER stress^8,9^. Therefore, abnormal amino acid substitutions of the ASD-related synaptic proteins are predisposed to accumulate in the ER and hamper their trafficking to the proper sites, attenuating innate synaptic functions.

Hevin, also known as secreted protein acidic and rich in cysteine-like 1 (Sparcl1), is a secreted protein. The C-terminus of Hevin has an amino acid sequence with approximately 60% identity to Sparc. It has been reported that Hevin and its family proteins are involved in metastasis, inflammation, angiogenesis, and apoptosis^10^. In the central nervous system, Hevin is expressed in astrocytes predominantly and is expressed in neurons to a lesser extent^11^. During brain development, Hevin is expressed in the radial glia and controls a detachment process that terminates neuronal migration in the pial surface of the cerebral cortex^12^. Hevin is also secreted from neural stem cells (NSCs) in the lateral ventricle subventricular zone (SVZ) and enhances glioma cell invasion^13^. Thus, it seems likely that Hevin is implicated in cell migration and invasion in a context-dependent manner. Importantly, it has been reported that Hevin reinforces the connection between pre- and post-synapses in the cerebral cortex^14^. On this occasion, Hevin interacts with both Neurexins1α (NRX1α) and NL1B and becomes a bridge between the thalamocortical synaptic connections in the cerebral cortex. Hevin has also been reported to enhance NMDAR-mediated synapse response^15^, demonstrating that one of the important functions of Hevin is to regulate synaptic connectivity and plasticity. Strikingly, genome-wide association studies have revealed that multiple mutations in *SPARCL1* gene increase the risk of ASD^5^, indicating that Hevin plays an important role in regulating synaptic connectivity and the brain environment.

A recent structural analysis reported how Hevin promotes the interaction between NRX and NL^16^. Hevin is composed of a flexible acidic region at the N-terminal region, follistatin-like (FS) domain at the center region, and extracellular calcium (EC) binding domain, including two EF-hand motifs (EF-hand 1: His^586^-Ala^618^ and EF-hand 2: His^625^-Phe^651^ amino acid in human, EF-hand 1: His^572^-Ala^604^ amino acid and EF-hand 2: His^611^-Phe^637^ amino acid in mouse) at the C-terminal region. Interestingly, the structure of the FS/EC region of Hevin differs from Sparc, although the Sparc shares high sequence identity with Hevin. The FS-domain is crucial for promoting the bridge between NRX and NL. Hevin and Sparc compete for interacting with NLs via FS-domain, thereby Sparc can antagonize the synaptogenic functions of Hevin. On the other hand, the EC-domain in both Hevin and Sparc interacts with extracellular matrix, such as Collagen I and V, in a Ca^2+^-dependent manner^16,17^. Therefore, analyzing functions of the EC domain is also of high importance to understand the full functions of Hevin.

In this study, we analyzed the importance of the EF-hand motif in Hevin and found that the integrity of EF-hand motif is necessary for proper trafficking to the extracellular space. In addition, a single amino acid substitution in the EF-hand motif, which is associated with ASD, causes Hevin to accumulate in the ER and activates the UPR pathway. These findings provide a possibility that protein accumulation by structural instability of Hevin contributes to the ASD pathogenesis.

## Results

### Hevin mutants lacking the EF-hand motif are generated in *Usp15* deficient brain

Previously, we reported that the deubiquitinating enzyme, USP15, deubiquitinates U6 snRNA-specific terminal uridylyl transferase 1 (TUT1) and regulates the spliceosome cascade^18^. A defect in *Usp15* increases the probability of splicing error and produces various abnormal variants. We also found that impaired USP15 induces the endoplasmic reticulum (ER) stress although the mechanism linking splicing error to ER stress has not been clarified^18^ (Fig.1A). Because *Usp15* gene is linked to ASD^2,19,20^, we looked for the ASD-associated abnormal variants, which have changed splicing upon *Usp15* deficiency and are involved in the UPR pathway. To do this, we checked the dataset of microarray analysis from our previous study^18^ and found that the 3’-region of hevin transcript tends to be lacking in *Usp15* deficient brains (Fig.1B, Supplementary Table). Thus, we speculated that the 3’-region of hevin transcript is susceptible to abnormalities in the spliceosome cascade regulated by USP15. We determined which base sequence of hevin transcript was altered in the *Usp15* deficient brain. To do this, we conducted a 3’-rapid amplification of cDNA ends (3’-RACE) and identified two 3’-fragments lacking the EF-hand motif. One was the EF-hand deletion mutant and another was the trans-splicing variant, which fused with a fragment of NADH:ubiquinone oxidoreductase subunit A11 (Ndufa11) (Fig. 1C to 1G). Because a truncated form of recombinant Hevin alters a protein structure^17^, we reasoned that these EF-hand deletion mutants show abnormal structure, resulting in a decrease of secretion efficiency.

**Figure 1.**
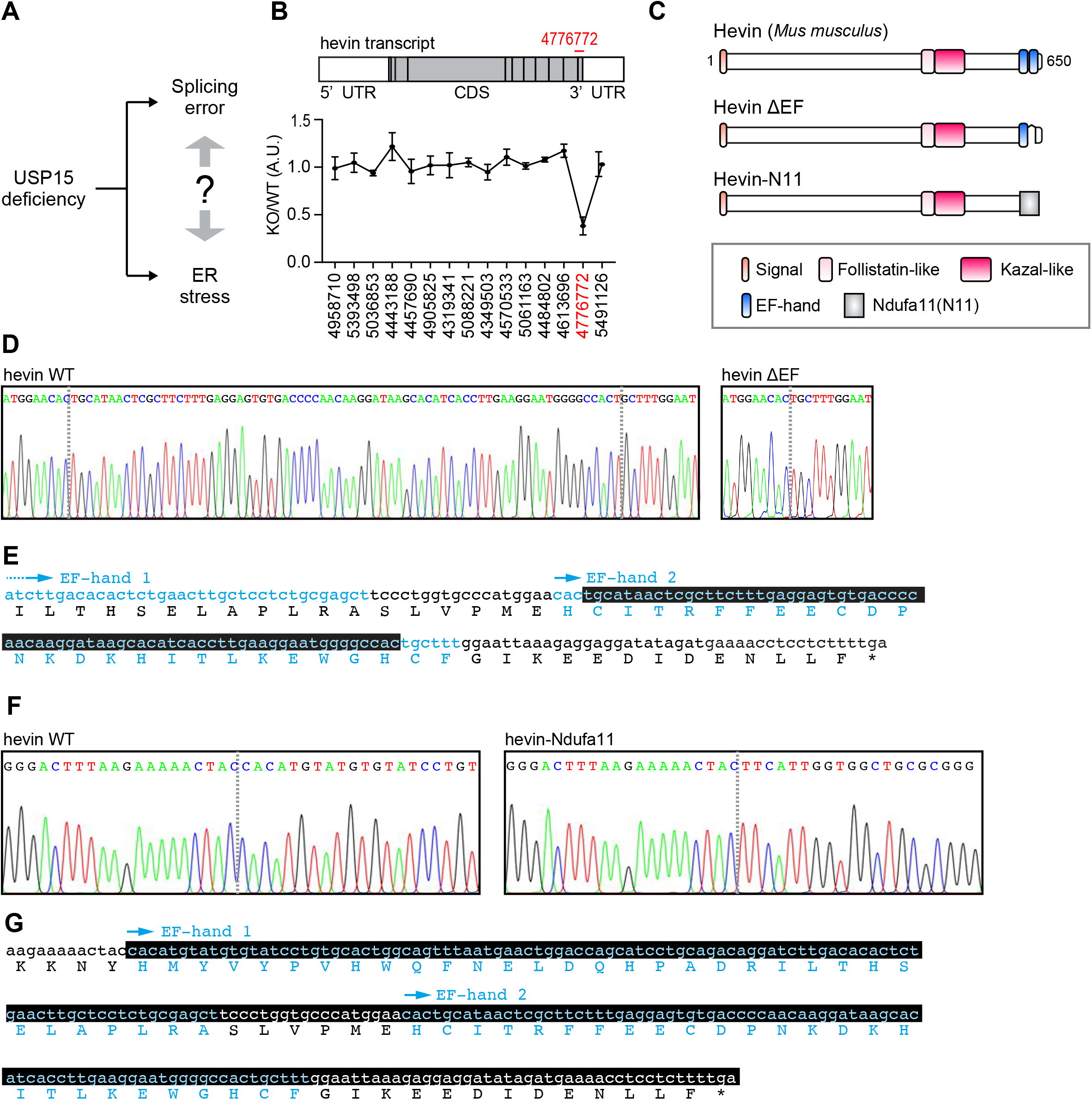
Hevin mutants lacking the EF-hand motif are generated in *Usp15* deficient brain. (A) The relationship between the splicing error and ER stress in *Usp15* deficient brain has not been clarified. (B) Schematic structure of hevin transcript in *Usp15* deficient mouse brain. The 3’-region of hevin transcript (PSR/Junction ID: 4776772) is prone to be missing in *Usp15* deficient brain. (C) The putative schematic structures of Hevin mutants. (D) to (G) Sequence analyses of the 3’-regions of hevin transcripts. (D) DNA sequencing chromatogram of 3’-region of hevin WT and ΔEF transcripts. (E) The 3’-region of hevin sequence. Black box indicates the lacking sequence in ΔEF mutant. Cyan indicates the EF-hand motifs. (F) DNA sequencing chromatogram of 3’-region of hevin WT and Ndufa11. (G) The 3’-region of hevin sequence. Black box indicates the lacking sequence in Hevin-N11 mutant. Cyan indicates the EF-hand motifs.

### Hevin mutants lacking the EF-hand motif activate the UPR signaling caused by abnormal trafficking

To investigate whether these mutants are secreted to the extracellular space, we constructed the expression plasmids of full-length Hevin mutants and transfected them into Neuro-2a cells. Although Hevin WT was efficiently secreted to the extracellular space, the deletion mutant of EF-hand 2 (Hevin ΔEF) and the fusion protein between Hevin lacking the EF-hand and Ndufa11 C-terminal region (Hevin-N11) showed reduced secretion efficiently compared to Hevin WT (Fig. 2A). These data suggest that the integrity of the EF-hand motifs is crucial for the secretion of Hevin. We next examined whether Hevin mutants accumulate in the ER. To do this, we introduced each plasmid into HeLa cells and stained using an anti-Hevin antibody. Consequently, Hevin mutants were prone to accumulate in the ER compared to Hevin WT (Fig. 2B and 2C). These data demonstrate that an abnormal EF-hand region suppresses a Hevin export from the ER. Next, we examined whether the accumulation of Hevin mutants activates the UPR pathway. Expression of both Hevin ΔEF and Hevin-N11 increased the mRNA of the UPR markers, *Bip* and *Chop* (Fig. 2D and 2E). Moreover, the protein level of BIP, which is the ER chaperone protein, was increased when Hevin mutants were expressed. Importantly, BIP expression level was similar to thapsigargin-stimulated cells (Fig. 2F), suggesting that Hevin EF-hand deletion mutants accumulate in the ER and activate the UPR pathway.

**Figure 2.**
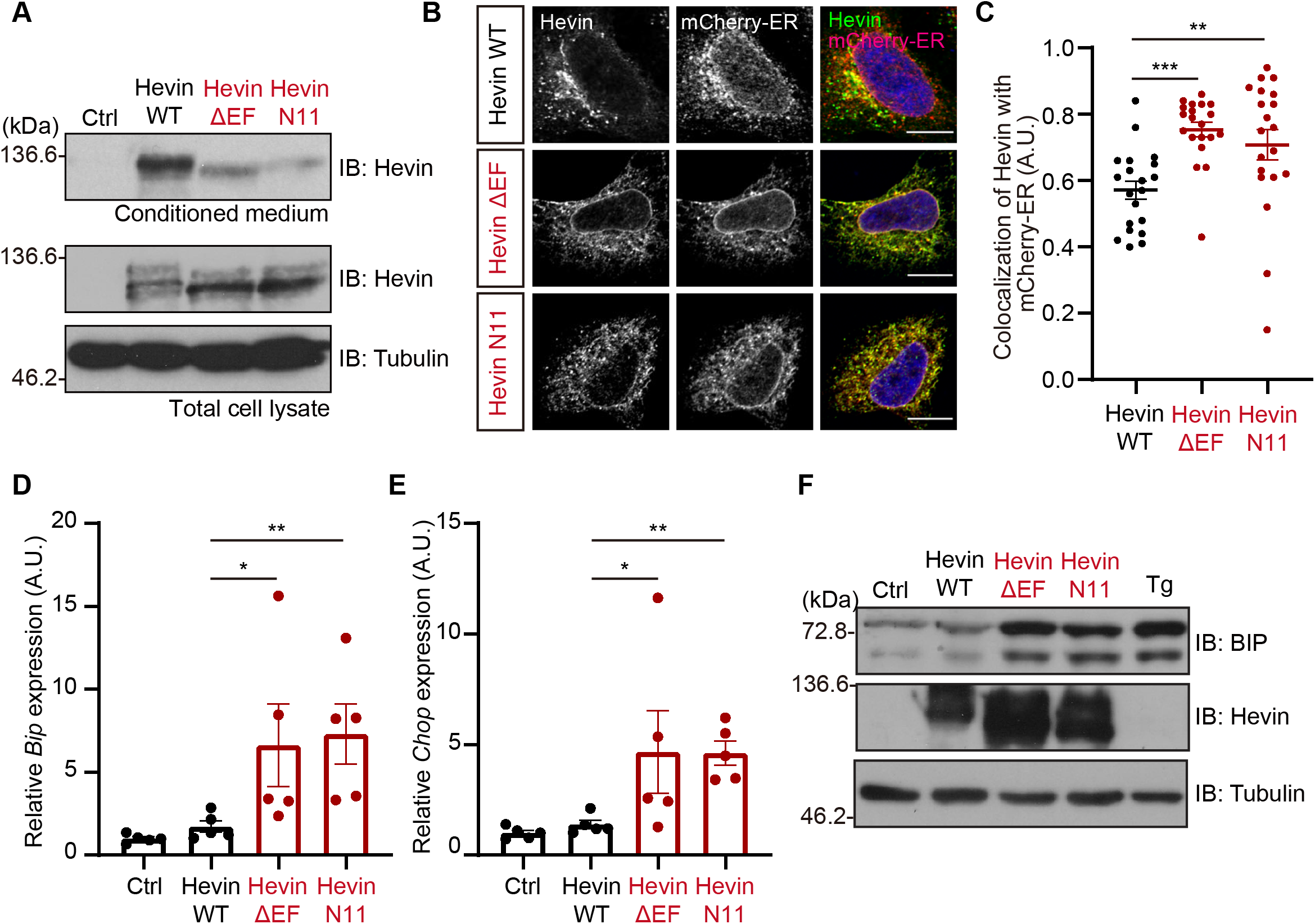
The Hevin mutants lacking the EF-hand motif activate the UPR signaling caused by abnormal trafficking. (A) Immunoblotting of Hevin WT and mutants (ΔEF and N11). Neuro-2a cells were transfected for 48 h with expression vectors for Hevin WT or mutants (ΔEF or N11). The cell lysates and conditioned medium were subjected to immunoblot analysis using anti-Hevin and Tubulin antibodies. (B) Immunostaining of Hevin and mutants (ΔEF and N11). HeLa cells were transfected for 24 h with expression vectors for Hevin and mCherry-ER and subjected to immunocytochemistry. Scale bar: 10 μm. (C) Quantification of Pearson’s correlation coefficient as the degree of colocalization in the panel B. n=20 cells, mean ± SEM, **P<0.01, ***P<0.001 versus Hevin WT, one-way analysis of variance (ANOVA) Dunnett’s test. (D)(E) Total RNAs isolated from the Hevin-overexpressed Neuro-2a cells were conducted with qPCR analysis for measurement of *Bip* and *Chop* mRNAs. 5S rRNA were used for normalization. n=5; mean ± SEM; *P<0.05, **P<0.01 versus Hevin WT, one-way ANOVA Dunnett’s test calculated using ΔCt value. (F) Immunoblotting of BIP, Hevin and Tubulin. HEK293T cells were transfected for 72 h with expression vectors for Hevin WT or mutants (ΔEF or N11). Cells were stimulated with 500 nM Thapsigarging (Tg) for 24 h. The cells were then subjected to immunoblot analysis using anti-BIP, Hevin and Tubulin antibodies.

### Hevin ASD-associated W647R mutation activates the UPR signaling

Both *SPARCL1* and *Usp15* mutations increase the risk of ASD. Importantly, previous transcriptome analyses have identified a variety of single amino acid substitutions in SPARCL1/Hevin that are linked to ASD. One of these mutants in which Trp^647^ is replaced with Arg is in the EF-hand motif that is also perturbed in *Usp15* knockouts (Fig. 3A, Supplementary Fig. 1). Strikingly, this amino acid is highly conserved among aves, rodents, and primates (Fig. 3B). Therefore, we postulated that this mutant [human W647R (hW647R)] could exhibit a similar cellular phenotype to the truncated Hevin mutants that we observed in *Usp15* KO mice in Fig. 2. First, to investigate whether this amino acid substitution in the EF-hand motif affects Hevin trafficking, we constructed the expression plasmid using the mouse Hevin sequence. Since human Trp^647^ corresponds to mouse Trp^633^ (Fig. 3B), we replaced mouse Trp^633^ with Arg [mouse W633R (mW633R)]. As we expected, the secretion efficiency in Hevin mW633R was attenuated compared to that in Hevin WT (Fig. 3C). In addition, Hevin mW633R mutant was mostly localized in the ER, as well as Hevin ΔEF and Hevin-N11 (Fig. 3D and 3E) and was not transported to the Golgi apparatus efficiently (Fig. 3F and 3G). Furthermore, expression of Hevin mW633R increased the mRNA level of the UPR marker, *Bip* and *Chop*, and protein level of BIP (Fig. 3H to 3J), suggesting that a single amino acid substitution in the EF-hand causes Hevin accumulation in the ER, followed by activation of the UPR signaling. Next, we investigated the mechanisms of how Hevin mutant increases ER stress. Since unfolded proteins promote dissociation of BIP from the ER stress sensors such as IRE1, leading to interacting with freed BIP, we confirmed whether Hevin mW633R acts as an unfolded protein-like behavior. Consequently, the binding affinity of Hevin mW633R to BIP is higher than that of WT (Fig. 3K and 3L). Importantly, Ca^2+^ existence has little effect on this binding. These data suggest that Hevin mutant activates the UPR signaling via increasing the binding affinity with BIP.

**Figure 3.**
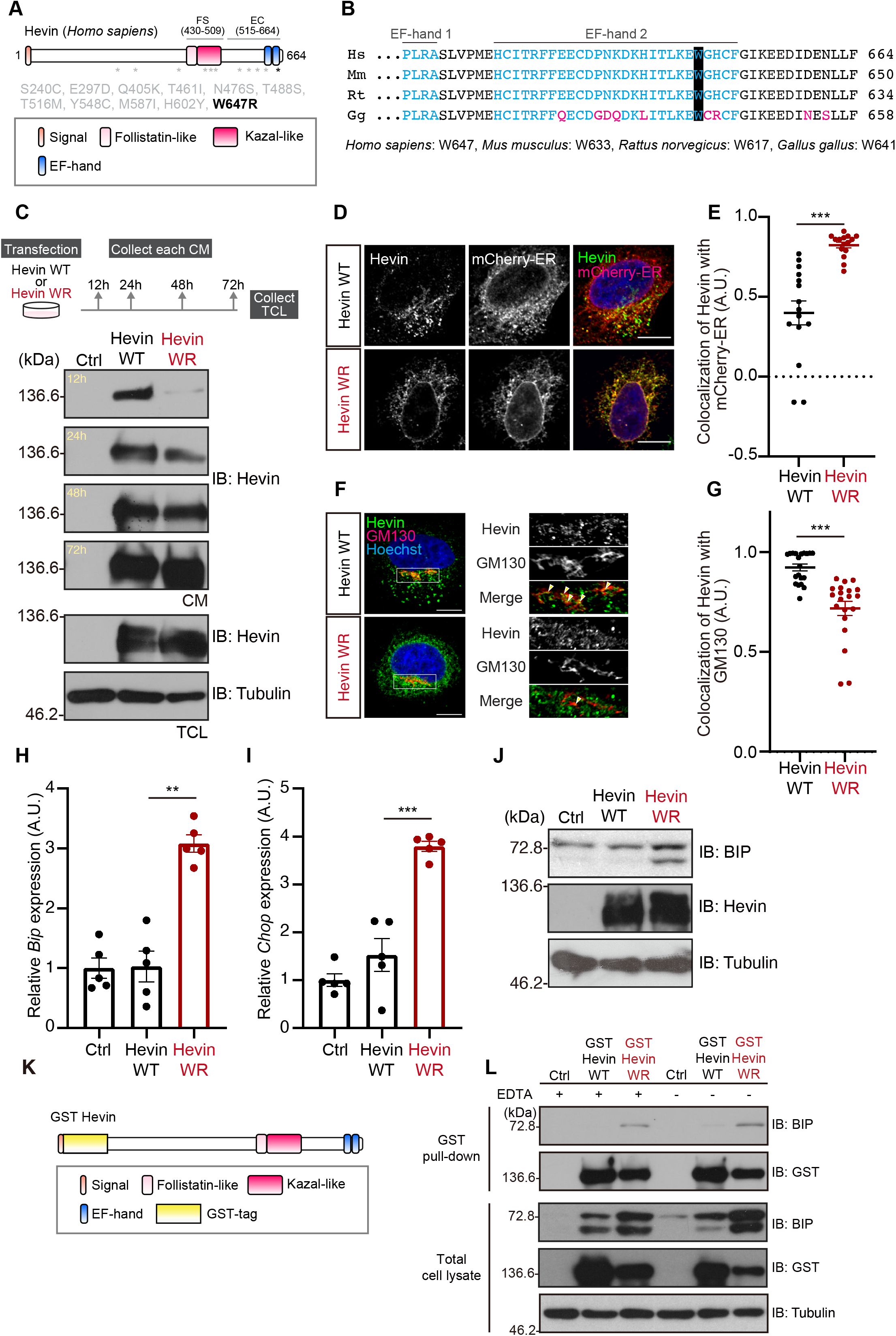
The ASD-associated mutant of Hevin activates the UPR signaling. (A) Schematic structure of ASD-associated Hevin mutants. (B) Amino acid of the EF-hand motifs in each species. The human Trp^647^ is highly conserved. Cyan indicates EF-hand motifs, Magenta indicates non-conserved amino acid. (C) Immunoblotting of Hevin WT and mW633R mutant. Neuro-2a cells were transfected with expression vectors and collected at the indicated times. The upper panel indicates the experimental schedule. The transfection efficiencies were confirmed by observing co-expressed EGFP fluorescence. The cell lysates and conditioned medium were subjected to immunoblot analysis using anti-Hevin and Tubulin antibodies. The secretion rate of WR mutant was delayed than that of WT. CM; conditioned medium, TCL; total cell lysate (D) Immunostaining of Hevin WT and mW633R mutant. HeLa cells were transfected for 24 hours with expression vectors for Hevin and mCherry-ER and subjected to immunocytochemistry. Scale bar: 10 μm. (E) Quantification of Pearson’s correlation coefficient as the degree of colocalization in the panel D. n=15 cells, mean ± SEM, ***P<0.001 by Student’s *t-*test. (F) Immunostaining of Hevin and mW633R mutant. HeLa cells were transfected for 24 h with expression vectors for Hevin and subjected to immunocytochemistry. Scale bar: 10 μm. (G) Quantification of Manders’ coefficient as the degree of colocalization of Hevin with GM130 in the panel F. n=20 cells, mean ± SEM, ***P<0.001 by Student’s *t-*test. (H)(I) Total RNAs isolated from the Hevin-overexpressed Neuro-2a cells were conducted with qPCR analysis for measurement of *Bip* and *Chop* mRNAs. 5S rRNA were used for normalization. n=5; **P<0.01, ***P<0.001 versus Hevin WT, one-way ANOVA Dunnett’s test calculated *p*-value using ΔCt value. (J) Immunoblotting of BIP, Hevin and Tubulin. Neuro-2a cells were transfected for 72 h with expression vectors for Hevin WT or mW633R mutants. The cell lysates were then subjected to immunoblot analysis using anti-BIP, Hevin and Tubulin antibodies. (K) Schematic structure of GST-Hevin. (L) Pull down of GST-Hevin and endogenous BIP in HEK293T cells. GST-Hevin was precipitated with Glutathione Sepharose beads and immunoblotted with anti-GST, BIP, and Tubulin antibodies.

### The structural destabilization of Hevin W647R mutant upon the hydrophobic core collapse

Because BIP tend to interacts with misfolded proteins in the ER^21^, we speculate that Hevin WR mutant exhibits structural instability. To verify this hypothesis, a set of μs-order all-atom molecular dynamics (MD) simulations was performed for the WT and hW647R of this protein in which the EC domain (515-664 amino acid) was analyzed (Fig. 4A). As the first analysis of both trajectories, the root-mean-square deviation (RMSD) for each initial structure during each MD simulation. As shown in the RMSD profiles, the hW647R system structurally fluctuated compared to the WT system since its RMSD value increased during the MD simulations, indicating that the structural destabilization of hW647R system is caused by a single amino acid substitution (Supplementary Fig. 2A to 2D). To address the origin of this structural destabilization, the solvent-accessible surface area (SASA) was measured in both systems. It is reasonable to measure the SASA values to address the mutational effect since the structural destabilization might be caused by a protein surface deformation, in which a part of protein surface of the WT system might be exposed to the external solvent upon this amino acid substitution. The SASA distributions of all the residues included within 8 Å around the mutation site were significantly different between the WT and hW647R systems (Fig. 4B and 4C). More specifically, the median value of SASA measured in the WT system was 2933 Å while that measured in the hW647R system was 3211 Å. These analyses indicate that this amino acid substitution causes the structural destabilization of the hW647R system. As the second analysis, the radial distribution function (RDF) of water was defined for all the residues included within 8 Å around the mutation site. The RDFs around each residue allow for identifications of essential residues that increase SASA. Among all the calculated distributions, the RDFs of water around V520 showed the characteristic profiles (Fig. 4D), where the RDF around V520 for the hW647R system significantly increased compared to that for the WT system. This increase of RDF for the hW647R system indicates that the inflow of water around V520 was induced by the amino acid substitution. Based on the difference in these RDFs, the magnitude of water inflow was compared among the corresponding residues. Quantitatively, the accumulation of the difference in RDFs around each residue between the WT and hW647R systems was measured for all the residues within 8 Å around the mutation site follows: Judging from the decomposed accumulation, the seven of the top ten residues with large RDF differences were hydrophobic residues (Fig. 4E). Therefore, the structural destabilization owing to this amino acid substitution might be attributed to the change in the inflow of water around these hydrophobic residues. Notably, the hydrophobic core formed in the WT system was collapsed in the hW647R system (Fig.4F and 4G). To evaluate the collapse of the hydrophobic core quantitively, the radius of gyration (Rg) of the seven hydrophobic residues was measured. The Rg value of these hydrophobic residues characterizes the size of each hydrophobic core. The distributions of Rg were different between both systems (Fig. 4H and 4I). More specifically, the median value of Rg measured in the WT system was 8.847 while that measured in the hW647R system was 9.675, indicating that the hydrophobic core of the hW647R system was collapsed by the inflow of water from the external region. The collapse of the native hydrophobic core is reasonable since the hydrophobic property of the Trp^647^ is lost by replacing it with Arg owing to this amino acid substitution. Therefore, it is likely that Hevin hW647R collapsed the hydrophobic core formed in the WT system, resulting in the structural destabilization of this protein.

**Figure 4.**
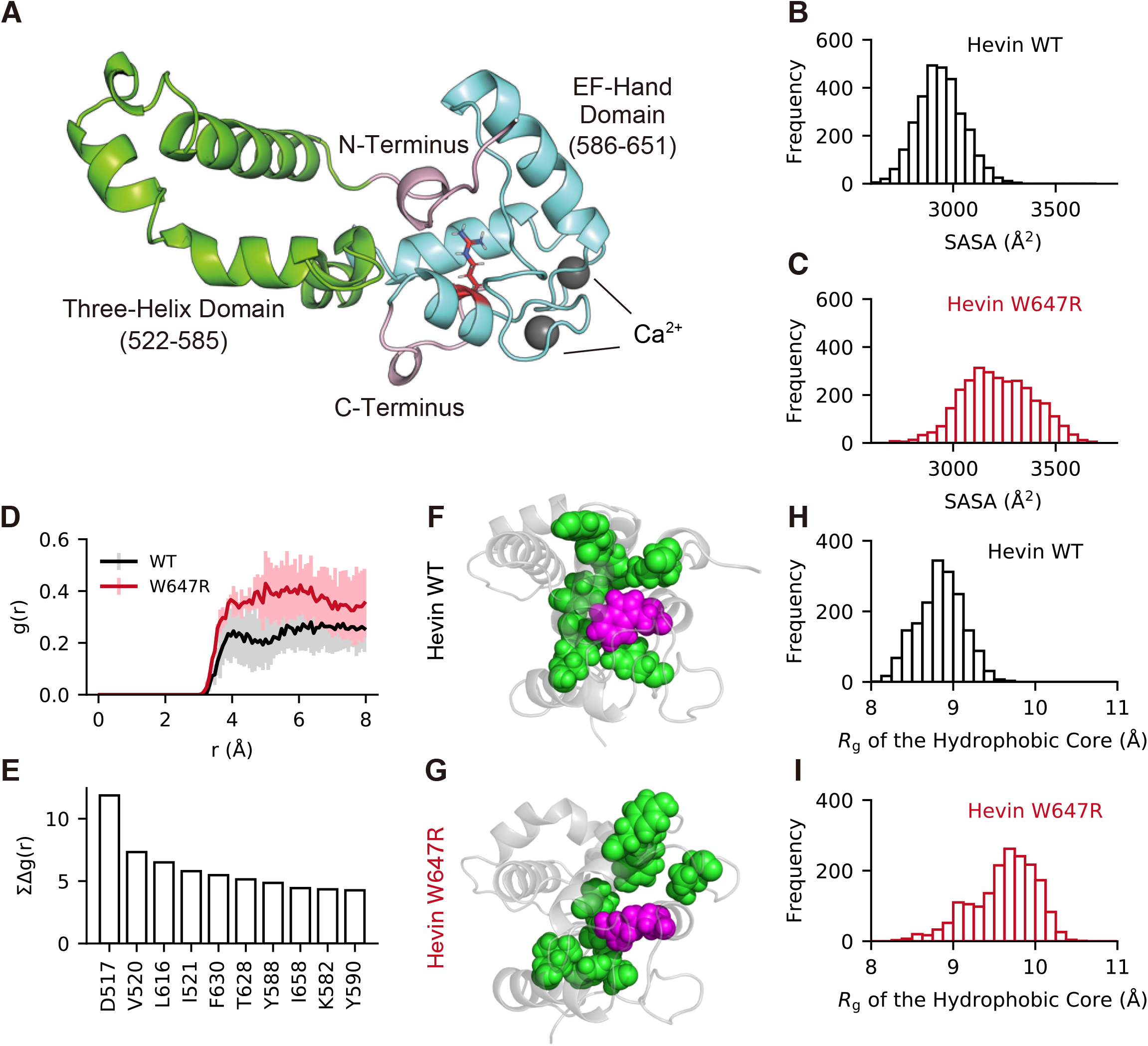
The structural destabilization of Hevin W647R mutant upon the hydrophobic core collapse. (A) The modeled structure of the WT Hevin system (EC domain). Each color represents the three-helix domain (green), EF-hand domain (cyan), and the N or C terminals (pink), respectively. The spheres (grey) represent Ca^2+^ ions. The structure is reconstituted from the data in the previous research^16^ (B)(C) The SASA distribution of all the residues included within 8 Å around the mutation site. (D) Characteristic RDFs of water around V520 with their standard deviations. (E) The accumulation of the difference in the RDFs of water between the WT and W647R systems, i.e., ∑_*r*_ *Δ*g(*r*) = ∑_*r*_ g(*r*)_WT_ − g(*r*)_W647R_, versus the top ten residues. (F)(G) A set of configurations of each hydrophobic core (D517, V520, L616, I521, F630, T628, Y588, I658, K582 and Y590), where the hydrophobic core and mutated residues are highlighted in green and magenta. (H, I) The Rg distributions of each hydrophobic core.

## Discussions

In this study, we identified that the defects in the Hevin EF-hand motif decrease secretion efficiency and accumulate in the ER. In addition, ASD-associated Hevin mutant also shows impaired trafficking and hamper an export from the ER. Indeed, Hevin mutant exposes a hydrophobic region, which is normally hidden inside, to the surface, inhibiting an export Hevin from the ER and activating the UPR pathway (Fig. 5). Thus, our findings provide a potential mechanism that links an ASD mutation in *SPARCL1* gene to UPR.

**Figure 5.**
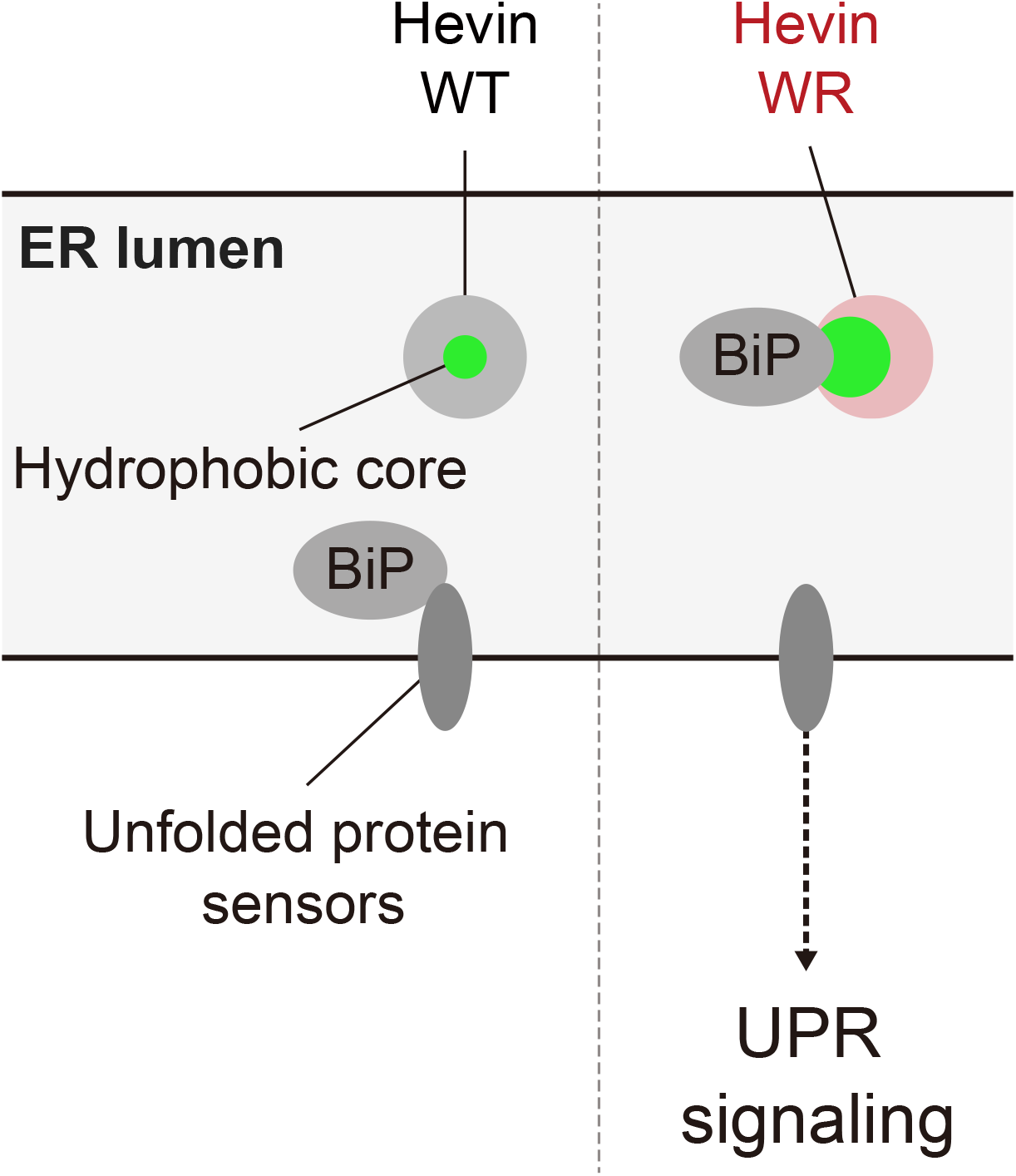
Model describing the dynamics of Hevin WT and WR mutant in the ER. BIP associates with Hevin WR mutant and triggers the UPR activation.

We found here that this single amino acid substitution in the EF-hand has two cellular phenotypes; First, the expression of Hevin WR activates the UPR pathway. Second, the mutation attenuates secretion efficiency. Thereby, we presume that the activation of the UPR pathway and/or a decrease of secretion efficiency are possible mechanisms that connect the Hevin dysfunctions to ASD. Previously, ER stress was proposed as a cellular mechanism related to the onset of ASD^22^. In fact, the UPR-related transcripts are dysregulated in the frontal cortices of ASD patients^23^. In most cases of regulating the UPR, three distinct pathways, inositol-requiring enzyme 1 (IRE1) - X-box binding protein 1 (XBP1) pathway, protein kinase R-like ER kinase (PERK) - eukaryotic initiation factor 2α (eIF2α) pathway and activating transcription factor 6 (ATF6) pathway are upregulated by the ER stress, such as abnormal Ca^2+^ homeostasis in the ER and perturbed the ER-Golgi trafficking. Because Hevin is predominantly expressed in astrocytes, it is likely that perturbed the UPR pathways in astrocytes could be an important mechanism underlying ASD pathogenesis. A tantalizing question for future research is the biological relevance of how ER stress response in astrocytes contribute to neural circuit and brain homeostasis. In *Caenorhabditis elegans*, overexpression of Xbp1 in astrocyte-like glial cells extends the life span via changing neuropeptide secretion machinery^24^. This observation demonstrates that upregulation of the UPR in astrocyte-like glial cells is pivotal for regulating ER stress resistance and longevity, exhibiting a positive effect in nematodes. Interestingly, a mutation in *Xbp1* is responsible for an onset of bipolar disorder^25^. Hence, these results raise the interesting hypothesis that dysfunctions of Xbp1 axis could also regulate not only neurodevelopmental but also psychiatric disorders. On the other hand, overactivation of PERK-eIF2α signaling in astrocytes causes neuronal loss and neurodegeneration in mice^26^. Strikingly, impaired UPR signaling markedly changes their secretory factors from astrocytes, suggesting that each UPR signaling in astrocyte regulates surrounding cells in a non-cell-autonomous manner. Thus, it is plausible that upregulation of ER stress induced by Hevin WR may change global secretome, resulting in circuit dysfunction that is causal to ASD. Another possibility is that the UPR signaling controls the timing of neural and glial differentiation. For instance, OASIS, a subfamily of ATF6, controls proper UPR signaling and regulates astrocyte differentiation in the embryonic stage^27^. Since perturbed glial differentiation arises developmental disorders, it is plausible that Hevin mutant-induced ER stress during the embryonic period influences glial differentiation, leading to a development of ASD. Previously, various maternal immune activations (MIA), such as exposures to environmental irritants and infections of pathogens, have been reported to cause ASD. Intriguingly, MIA activates the PERKs-eIF2α pathway and inhibits new protein synthesis. Either genetic or pharmacological inhibition of this pathway improves MIA-induced ASD-like behavior^28^, supporting the idea that ER stress is related to ASD.

Hevin promotes synaptogenesis through stabilizing the interaction between NRX1α and NL1B^14^. The structure of Hevin is divided into mainly three parts, the acidic domain, FS domain, and EC domain^16^. The FS domain binds to NRX1α and NL1B concurrently, and the EC domain, of Hevin including the EF-hand motif, is predicted to be essential for interacting with the collagen in a Ca^2+^-dependent manner^16,17^. We found that only a small amount of Hevin mW633R mutant can be secreted to the extracellular space even in a mutation of the EF-hand. These data imply that exposure to the hydrophobic region of the C-terminus may acquire novel functions that disturb innate functions in the extracellular space even in the existence of Ca^2+^. One hypothesis is that the binding affinity between Hevin and Collagen proteins is changed and affects functions of synaptogenesis. Ehlers-Danlos syndromes, which exhibit autism-like phenotypes arise from mutation in *Col1A* and *Col5A* genes^29^. Therefore, the Hevin WR mutant could also impact the conditions of the extracellular matrix and brain microenvironment. Previous studies have reported that Sparc, a homolog of Hevin, suppresses synaptogenesis. Intriguingly, the deletion mutant lacking an acidic domain (SLF) exhibits a similar phenotype of Sparc in which SLF suppresses synaptogenesis^30^. Hevin is also cleaved at the center region (approximately 350 amino acids) by a metalloprotease, ADAMTS4, and is produced the fragment containing both FS and EC domains^31^. Therefore, it is likely that Hevin hW647R mutant acquires a distinct function as well as Sparc and SLF. However, further studies will be needed to clarify this possibility in the future.

Previous studies have reported that other mutations in *SPARCL1 gene* associated with ASD have been revealed, and most mutations reside in the FS domain and EC domain^5^ (see Fig. 3A). Mutations in which Thr^516^ and Met^587^ are replaced with Met and Ile respectively are next to the residue (Asp^517^ and Tyr^588^) with large RDF differences. Thus, it is likely that the native hydrophobic core is collapsed when these amino acids are substituted as well as WR mutant. ASD-associated amino acid substitutions around the EF-hand motif may change the native hydrophobic core, resulting in the structural destabilization of Hevin. Notably, Ca^2+^ deprivation has less effect on the binding between BIP and Hevin WR mutant. Thus, it is plausible that exposure of hydrophobic region covered by not only Trp^647^ but also Thr^516^ and Met^587^ impairs Hevin secretion and synaptogenesis irrespectively of Ca^2+^.

In summary, here we found that the EF-hand motif in Hevin plays an important for proper trafficking to the extracellular spaces. A loss of this domain causes protein accumulation in the ER and activates the UPR pathway. Furthermore, we found that ASD-associated mutation in which Trp^647^ is replaced with Arg exhibits a similar phenotype in cells. Importantly, in hW647R mutant Hevin the hydrophobic core of the EF hand is exposed to the surface likely causing structural instability. Taken together, our findings provide multiple molecular mechanisms linking an ASD point mutation in *SPARCL1* gene with cellular phenotypes underlying the onset of ASD.

## Materials and Methods

### Plasmids

The mouse Hevin cDNA was amplified from pMD-mSPARCL1 (Sino Biological Inc. MG50544-M) and cloned into pCRblunt. The deletion mutant of EF-hand (Hevin ΔEF) was constructed using the inverse PCR method. Briefly, Hevin ΔEF cDNA was amplified from pCRblunt-Hevin, followed by conducting a T4 polynucleotide kinase reaction (TAKARA). After phosphorylation, this fragment was self-ligated and constructed a circular plasmid as a pCRb-Hevin ΔEF. The Hevin-Ndufa11 (Hevin-N11) fragment was amplified from Hevin cDNA and Ndufa11 cDNA obtained from 3’-RACE method, followed by ligating these fragments using an In-Fusion system (TAKARA). Hevin-N11 fragment was inserted into pCR-blunt using In-Fusion system. The gene encoding Hevin mW633R was generated by a quick-change method using KOD polymerase. The GST-Hevin proteins are generated by the SLiCE method^32^. Briefly, the full-length Hevin cDNA was amplified using KOD plus DNA polymerase (TOYOBO) and inserted into the BglII sites of pCS4 vector. The GST-tagged Hevin and Hevin mW633R mutant were generated using SLiCE reaction. The GST fragment is subcloned from pGEX-6P-1 plasmid into the site between Hevin signal-peptide and another part using SLiCE reaction. The primers used were as follows:

Full-length Hevin forward, 5’-aaagatctgccaccatgaaggctgtgcttctcc-3’;

Full-length Hevin reverse, 5’-ggagatcttcaaaagaggaggttttcatctatatcctcctc-3’;

Hevin ΔEF inverse sense, 5’-tgctttggaattaaagaggaggatatagatg-3’;

Hevin ΔEF inverse antisense, 5’-gtgttccatgggcaccaggg-3’;

N-terminal Hevin forward, 5’-gctggaattcaggagtgaaggctgtgcttc-3’;

N-terminal Hevin reverse, 5’-gatggtctccagccaagagtctcttctcatcc-3’;

C-terminal Hevin-Ndufa11 mutants forward, 5’-gagactcttggctggagaccatcccattg-3’;

C-terminal Hevin-Ndufa11 mutants reverse, 5’-tctgcagaattcaggacaccttgggggtag-3’;

Hevin W633R mutant forward, 5’-gaaggaacggggccactgc-3’;

Hevin W633R mutant reverse, 5’-gcagtggccccgttccttc-3’;

Hevin (pCS4) for SLiCE forward, 5’-tcgtgacatcccgacaagtacaaggtttctctc-3’;

Hevin (pCS4) for SLiCE reverse, 5’-tataggggatgccacagcggttcccaag-3’;

GST for SLiCE forward, 5’-ctgtggcatcccctatactaggttattgg-3’;

GST for SLiCE reverse, 5’-ttgtcgggatgtcacgatgcgg-3’;

### 3’-Rapid amplification of cDNA ends (3’-RACE)

Total RNAs from wild-type and *USP15* deficient brain were isolated by ISOGEN II (NIPPON GENE) according to the manufacturer’s instructions. The cDNAs were synthesized by reverse transcriptase and 100 units ReverTra Ace (TOYOBO) together with 25 pmol random hexamer primer (TOYOBO), 20 nmol dNTPs and total RNAs. The 3’-region of hevin transcript was determined using 3’-RACE method. The first-strand synthesis was conducted using the Hevin outside primer and Hevin adaptor primer. Then, the second round of PCR was conducted using the Hevin inside primer and Hevin amplification primer. The 3’-fragment amplified by 3’-RACE was inserted into pCR-blunt and analyze these DNA sequences. The primers used were as follows:

Hevin outside primer, 5’-aaatgctgaaccttcagatgagggc-3’;

Hevin adaptor primer, 5’-ggccacgcgtcgactagtacttttttttttttttttt-3’;

Hevin inside primer, 5’-agagactcttggctggagaccatc-3’;

Hevin amplification primer, 5’-ggccacgcgtcgactagtac-3’;

### Antibodies

For immunoblotting goat anti-Hevin (1:1000, R&D Systems, Cat# AF2836), mouse anti-Tubulin (1:1000, Sigma, DM1A), rabbit anti-BIP (1:1000, Cell Signaling, CB0B12), mouse anti-GST (1:500, Santa Cruz, B14) antibodies were used as primary antibodies. The horseradish peroxidase-conjugated anti-mouse IgG, anti-rabbit IgG, and anti-goat IgG (1:20000 respectively, SeraCare) antibodies were used as secondary antibodies. For immunofluorescence, rabbit anti-RFP (1:500, Rockland, Cat# 600-401-379), goat anti-Hevin (1:500, R&D Systems, Cat# AF2836), and mouse anti-GM130 (1:500, MBL, Cat# M179-3) were used as primary antibodies. The Alexa Fluor 488 donkey anti-goat IgG H&L (1:500, Thermo Fisher Scientific), DyLight 594 donkey anti-rabbit IgG H&L (1:500, abcam) and DyLight 594 donkey anti-mouse IgG H&L (1:500, abcam) antibodies were used as secondary antibodies.

### Cell culture and transfection

HEK293T cells and HeLa cells were cultured in Dulbecco’s modified Eagle’s medium (High glucose) (Wako) containing 5% fetal bovine serum (FBS), 100 units penicillin and 100 mg streptomycin (P/S Thermo Fisher Scientific). Cells were transfected using polyethyleneimine MAX (PEI Max) (Polyscience). The amount of the plasmids and PEI Max were optimized in proportion to the relative surface area and the number of cells. Cells were plated 1.5 – 5.0 × 10^5^ cells in 4 ml per 60 mm-dish and incubated at 5% CO2 and 37 °C for 1 d. The plasmids (8.0 to 16.0 μg) were mixed with 500 μl of Opti-MEM (Thermo Fisher) and 1.0 μg/μl of PEI Max (32.0 to 64.0 μl) was mixed with 500 μl of Opti-MEM in another tube. Both solutions were combined and incubated for 20 min at room temperature, followed by adding these mixtures to cells. After incubation for 1 h, the media were changed to fresh culture medium. Neuro-2a cells were cultured in Eagle’s minimum essential medium (Wako) containing 10% FBS and P/S. Cells were plated 1.5 × 10^5^ cells on 6 well plate and incubated at 5% CO2 and 37°C for 1 d. The plasmids (4.0 μg) were mixed with 250 μl of Opti-MEM (Thermo Fisher) and 1.0 μg/μl of PEI Max (8.0 μl) was mixed with 250 μl of Opti-MEM in another tube. Both solutions were combined and incubated for 20 min at room temperature, followed by adding these mixtures to cells. After incubation for 4 h, the media were changed to fresh culture medium.

### Immunocytochemistry

HeLa cells were plated at 5.0 × 10^4^ cells and incubated on 15 mm coverslips in 12-well plate for 1 d and then transfected with indicated plasmids. After the incubation for 24 h, the cells were washed in PBS and fixed with 4% paraformaldehyde (Merck KGaA) in PBS for 10 min at 4°C. The coverslips were washed in PBS and blocked with 5% bovine serum albumin (BSA; Wako) in PBS with 0.4% Triton X-100 (MP Biomedicals), then incubated with primary antibodies diluted in blocking solution for overnight at 4°C. After washing with PBS, the cells were incubated with secondary antibodies diluted in blocking solution for 30 min at room temperature. Nuclei were stained with 10 μg/ml Hoechst 33342 (Life Technologies). The coverslips were then mounted onto slides using FLUOROSHIELD Mounting Medium (ImmunoBioScience). Fluorescent images were obtained using a confocal laser scanning fluorescence microscopy (ZEISS, LSM700) with 63× (Plan-Apochromat 63×/1.4 Oil DIC) or 100× (αPlan-Apochromat 100×/1.46 Oil DIC) objective. The diode excitation lasers (Diode 405, Diode 488, and Diode 555) were operated and directed to a photomultiplier tube (LSM T-PMT, Carl Zeiss) through a series of band pass filters (Ch1:BP420-475 + BP500-610, Ch2:BP490-635, and Ch3:BP585-1000).

### Colocalization analysis

Colocalization analysis of immunofluorescence signals was performed according to the previous study^33^. The Colocalization values (Pearson’s co-efficients or Manders’ co-efficients) were calculated using immunofluorescence signals of Hevin and either mCherry-ER or GM130 from representative images by Fiji plugin Coloc-2. All data were reproduced in at least two independent experiments.

### GST Pull down assay

HEK293T cells were plated at 5.0 × 10^5^ cells and incubated for 1 d and then transfected with indicated plasmids. After the incubation for 48 h, cells were washed with PBS and collected with lysis buffer [20 mM Tris-HCl (pH 7.5), 150 mM NaCl, 0.5% NP-40, 1 mM DTT, 1 mM PMSF, 3 μg/ml leupeptin, 3 μg/ml Pepstatin A, and 5 μg/ml aprotinin] in the presence or absence of 1 mM EDTA. Cell lysates were centrifuged at 14,000 rpm for 5 min. The supernatants were mixed with Glutathione Sepharose beads (GE Helthcare) and were incubated for 2 h at 4 °C with rotation. The precipitants were washed 3 times with lysis buffer and subjected to immunoblot analysis. All data were reproduced in at least two independent experiments.

### Immunoblot analysis

HEK293T cells or Neuro-2a cells were plated at 0.8 or 1.5 × 10^5^ cells on six-well plate and incubated at 37°C with 5% CO2 for 1 d. Cells were transfected and then incubated for 3 d. The cells were collected with lysis buffer [20 mM Tris-HCl (pH 7.5), 150 mM NaCl, 1 mM EDTA, 0.5% NP-40, and 1 mM DTT]. Cell lysates were centrifuged at 14,000 rpm for 5 min. The supernatant was run on SDS-PAGE for protein separation, followed by electrophoretic transfer to a polyvinylidene difluoride membrane (Pall). After 1 h blocking by 5% skim milk at room temperature, membranes were incubated with primary antibodies overnight at 4°C. The proteins on membrane were then detected with HRP-conjugated secondary antibodies and chemiluminescence reagents (Chemi-Lumi One Super, Nacarai tesque). All data were reproduced in at least two independent experiments.

### Quantitative real-time PCR (qPCR)

Neuro-2a cells (1.5 × 10^5^ cells/ six-well plate) were cultured for 24 h and then transfected with indicated plasmids. After the incubation for 48 h, the cells were washed in PBS and collected with ISOGEN II (NIPPON GENE). Total RNAs from cells were isolated according to the ISOGEN II manufacturer’s instructions. The cDNAs were synthesized by reverse transcriptase, 100 units ReverTra Ace (TOYOBO) together with 25 pmol Random Primer (nonamer; TOYOBO), 20 nmol dNTPs and 500 ng total RNAs. The qPCR was performed in 96-well plate (Scientific Specialties) using THUNDERBIRD SYBR qPCR Mix (TOYOBO) in Applied Biosystems 7900HT Fast Real Time PCR System (Applied Biosystems). The relative quantity of the target expression was calculated by 2^-ΔΔCt^ methods using SDS Software 2.4.2 (Applied Biosystems) with the following calculation. The relative quantity = 2^-ΔΔCt^, ΔΔCt = (Ct^target^ – Ct^5S^)sample – (Ct^target^ – Ct^5S^)reference; Ct, threshold cycle. All data were reproduced in at least two independent experiments. The primers used were as follows:

5S rRNA forward, 5’-cggccataccaccctgaac-3’;

5S rRNA reverse, 5’-gcggtctcccatccaagtac-3’;

*Bip* forward, 5’-gagactgctgaggcgtattt-3’;

*Bip* reverse, 5’-cctcatgacattcagtccag-3’;

*Chop* forward, 5’-ctggaagcctggtatgaggat-3’;

*Chop* reverse, 5’-cagggtcaagagtagtgaaggt-3’;

### Molecular dynamics (MD) simulation

The WT and W647R systems of Hevin were prepared to conduct MD simulation. In each system, the EC domain (511-663) of Hevin was only considered. The WT and mutant (W647R) systems were modeled using a crystal structure (PDB code: 7KBU)^16^. To model the W647R system, this amino acid substitution was induced using the Pymol program (The PyMOL Molecular Graphics System, Version 2.0 Schrödinger, LLC.). In each system, the missing amino-acid residues were added using the SWISS-MODEL program^34^. The modeled structures were solvated with the TIP3P water model^35^ in an orthorhombic box. The LINCS^36^ and SETTLE^37^ algorithms were adopted to extend the MD time step to 2 fs. The modified Berendsen thermostat^38^ and Parrinello–Rahman method^39,40^ controlled the temperatures and pressures of all the systems, respectively. The particle mesh Ewald method^41^ evaluated electrostatic interactions using a real-space cutoff of 10 Å. The cutoff value for the van der Waals interactions was set to 10 Å. All the MD simulations were performed with the GPU version of the GROMACS 2020 package (Abraham DvdS, M.J., Lindahl, E., Hess, B., and the GROMACS development team. GROMACS User Manual version 2019) using the Amber 14SBonlysc force field^42^. Before conformational sampling, energy minimizations on the initially modeled systems removed the steric crashes of atoms. Subsequently, 100-ps *NVT* (*T* = 300 K) and 100-ps *NPT* (*T* = 300 K and P = 1 bar) MD simulations relaxed each system. Finally, the final snapshots of the *NPT* simulations on each system were specified as the starting configurations of their production runs. To generate statistically reliable MD trajectories, multiple 1-μs MD simulations (1-μs × 3 runs for each system) were independently started from each relaxed configuration by regenerating their initial velocities. Finally, the first 0.4-μs trajectories were considered as the equilibration phase, and the remaining 0.6-μs trajectories were used for analyses in each system.

### Statistical analysis

Prism ver.8.4.3 software (GraphPad Software, Inc.) was used for all statistical analyses. Statistical significance was analyzed using Student’s *t*-test and ANOVA followed by Dunnett’s test.

## Acknowledgements

We thank the lab members for helpful discussions and technical support. MD simulations were partially performed with a supercomputer (Cygnus) provided by the Multidisciplinary Cooperative Research Program (MCRP) in the Center for Computational Sciences (CCS), University of Tsukuba, Japan (project code: LSC and MOLBIO). This work was supported by Grant-in Aid from the Ministry of Education, Science, Sports and Culture of Japan JSPS KAKENHI (16K07339, 16KK0158, 20K05951), Astellas Foundation for Research on Metabolic Disorders and Gout and uric acid foundation of Japan to FT. CE is an HHMI Investigator.

## Author contributions

T.T. and F.T. designed research; T.T., T.Y., R.M., J.K. and F.T. performed experiments; T.T., T.Y., R.M., J.K., R.H., and F.T. analyzed data; T.T., T.Y., R.H., C.E. and F.T. wrote the paper. Study supervision: Y.S., C.E. and F.T

## Additional information

Data availability. The GEO accession number for the array data set is GSE145385^18^. The PDB code of Hevin crystal structure is 7KBU^16^. The authors declare no competing interests.

