## Supplementary Fig. for "Autism-associated mutation in Hevin/Sparcl1 induces endoplasmic reticulum stress through structural instability"

```
   1 atgaagactgggctttttttcctatgtctcttgggaactgcagctgcaatcccgacaaatgcaagattattatctgatcattccaaaccaactgctgaaacggtagcacctgacaacact
   1  M  K  T  G  L  F  F  L  C  L  L  G  T  A  A  A  I  P  T  N  A  R  L  L  S  D  H  S  K  P  T  A  E  T  V  A  P  D  N  T

 121 gcaatccccagtttaagggctgaagctgaagaaaatgaaaaagaaacagcagtatccacagaagacgattcccaccataaggctgaaaaatcatcagtactaaagtcaaaagaggaaagc
  41  A  I  P  S  L  R  A  E  A  E  E  N  E  K  E  T  A  V  S  T  E  D  D  S  H  H  K  A  E  K  S  S  V  L  K  S  K  E  E  S

 241 catgaacagtcagcagaacagggcaagagttctagccaagagctgggattgaaggatcaagaggacagtgatggtcacttaagtgtgaatttggagtatgcaccaactgaaggtacattg
  81  H  E  Q  S  A  E  Q  G  K  S  S  S  Q  E  L  G  L  K  D  Q  E  D  S  D  G  H  L  S  V  N  L  E  Y  A  P  T  E  G  T  L

 361 gacataaaagaagatatgagtgagcctcaggagaaaaaactctcagagaacactgatttttttggctcctggtgttagttccttcacagattctaaccaacaagaaagtatcacaaagaga
 121  D  I  K  E  D  M  S  E  P  Q  E  K  K  L  S  E  N  T  D  F  L  A  P  G  V  S  S  F  T  D  S  N  Q  Q  E  S  I  T  K  R
   1                 M  S  E  P  Q  E  K  K  L  S  E  N  T  D  F  L  A  P  G  V  S  S  F  T  D  S  N  Q  Q  E  S  I  T  K  R

 481 gaggaaaaccaagaacaacctagaaattattcacatcatcagttgaacaggagcagtaaacatagccaaggcctaagggatcaaggaaaccaagagcaggatccaaatatttccaatgga
 161  E  E  N  Q  E  Q  P  R  N  Y  S  H  H  Q  L  N  R  S  S  K  H  S  Q  G  L  R  D  Q  G  N  Q  E  Q  D  P  N  I  S  N  G
  36  E  E  N  Q  E  Q  P  R  N  Y  S  H  H  Q  L  N  R  S  S  K  H  S  Q  G  L  R  D  Q  G  N  Q  E  Q  D  P  N  I  S  N  G

 601 gaagaggaagaagaaaaagagccaggtgaagttggtacccacaatgataaccaagaaagaaagacagaattgcccagggagcatgctaacagcaagcaggaggaagacaatacccaatct
 201  E  E  E  E  E  K  E  P  G  E  V  G  T  H  N  D  N  Q  E  R  K  T  E  L  P  R  E  H  A  N  S  K  Q  E  E  D  N  T  Q  S
  76  E  E  E  E  E  K  E  P  G  E  V  G  T  H  N  D  N  Q  E  R  K  T  E  L  P  R  E  H  A  N  S  K  Q  E  E  D  N  T  Q  S

 721 gatgatattttggaagagtctgatcaaccaactcaagtaagcaagatgcaggaggatgaatttgatcagggtaaccaagaacaagaagataactccaatgcagaaatggaagaggaaaat
 241  D  D  I  L  E  E  S  D  Q  P  T  Q  V  S  K  M  Q  E  D  E  F  D  Q  G  N  Q  E  Q  E  D  N  S  N  A  E  M  E  E  E  N
 116  D  D  I  L  E  E  S  D  Q  P  T  Q  V  S  K  M  Q  E  D  E  F  D  Q  G  N  Q  E  Q  E  D  N  S  N  A  E  M  E  E  E  N

 841 gcatcgaacgtcaataagcacattcaagaaactgaatggcagagtcaagagggtaaaactggcctagaagctatcagcaaccacaaagagacagaagaaaagactgtttctgaggctctg
 281  A  S  N  V  N  K  H  I  Q  E  T  E  W  Q  S  Q  E  G  K  T  G  L  E  A  I  S  N  H  K  E  T  E  E  K  T  V  S  E  A  L
 156  A  S  N  V  N  K  H  I  Q  E  T  E  W  Q  S  Q  E  G  K  T  G  L  E  A  I  S  N  H  K  E  T  E  E  K  T  V  S  E  A  L

 961 ctcatggaacctactgatgatggtaataccacgcccagaaatcatggagttgatgatgatggcgatgatgatggcgatgatggcggcactgatggccccaggcacagtgcaagtgatgac
 321  L  M  E  P  T  D  D  G  N  T  T  P  R  N  H  G  V  D  D  D  G  D  D  D  G  D  D  G  G  T  D  G  P  R  H  S  A  S  D  D
 196  L  M  E  P  T  D  D  G  N  T  T  P  R  N  H  G  V  D  D  D  G  D  D  D  G  D  D  G  G  T  D  G  P  R  H  S  A  S  D  D

1081 tacttcatcccaagccaggcctttctggaggccgagagagctcaatccattgcctatcacctcaaaattgaggagcaaagagaaaaagtacatgaaaatgaaaatataggtaccactgag
 361  Y  F  I  P  S  Q  A  F  L  E  A  E  R  A  Q  S  I  A  Y  H  L  K  I  E  E  Q  R  E  K  V  H  E  N  E  N  I  G  T  T  E
 236  Y  F  I  P  S  Q  A  F  L  E  A  E  R  A  Q  S  I  A  Y  H  L  K  I  E  E  Q  R  E  K  V  H  E  N  E  N  I  G  T  T  E

1201 cctggagagcaccaagaggccaagaaagcagagaactcatcaaatgaggaggaaacgtcaagtgaaggcaacatgagggtgcatgctgtggattcttgcatgagcttccagtgtaaaaga
 401  P  G  E  H  Q  E  A  K  K  A  E  N  S  S  N  E  E  E  T  S  S  E  G  N  M  R  V  H  A  V  D  S  C  M  S  F  Q  C  K  R
 276  P  G  E  H  Q  E  A  K  K  A  E  N  S  S  N  E  E  E  T  S  S  E  G  N  M  R  V  H  A  V  D  S  C  M  S  F  Q  C  K  R
                                                                                            FS domain

1321 ggccacatctgtaaggcagaccaacagggaaaacctcactgtgtctgccaggatccagtgacttgtcctccaacaaaaccccttgatcaagtttgtggcactgacaatcagacctatgct
 441  G  H  I  C  K  A  D  Q  Q  G  K  P  H  C  V  C  Q  D  P  V  T  C  P  P  T  K  P  L  D  Q  V  C  G  T  D  N  Q  T  Y  A
 316  G  H  I  C  K  A  D  Q  Q  G  K  P  H  C  V  C  Q  D  P  V  T  C  P  P  T  K  P  L  D  Q  V  C  G  T  D  N  Q  T  Y  A

1441 agttcctgtcatctattcgctactaaatgcagactggaggggaccaaaaaggggcatcaactccagctggattattttggagcctgcaaatctattcctacttgtacggactttgaagtg
 481  S  S  C  H  L  F  A  T  K  C  R  L  E  G  T  K  K  G  H  Q  L  Q  L  D  Y  F  G  A  C  K  S  I  P  T  C  T  D  F  E  V
 356  S  S  C  H  L  F  A  T  K  C  R  L  E  G  T  K  K  G  H  Q  L  Q  L  D  Y  F  G  A  C  K  S  I  P  T  C  T  D  F  E  V
                                                                                                             EC domain

1561 attcagtttcctctacggatgagagactggctcaagaatatcctcatgcagctttatgaagccaactctgaacacgctggttatctaaatgagaagcagagaaataaagtcaagaaaatt
 521  I  Q  F  P  L  R  M  R  D  W  L  K  N  I  L  M  Q  L  Y  E  A  N  S  E  H  A  G  Y  L  N  E  K  Q  R  N  K  V  K  K  I
 396  I  Q  F  P  L  R  M  R  D  W  L  K  N  I  L  M  Q  L  Y  E  A  N  S  E  H  A  G  Y  L  N  E  K  Q  R  N  K  V  K  K  I

1681 tacctggatgaaaagaggcttttggctggggaccatcccattgatcttctcttaagggactttaagaaaaactaccacatgtatgtgtatcctgtgcactggcagtttagtgaacttgac
 561  Y  L  D  E  K  R  L  L  A  G  D  H  P  I  D  L  L  L  R  D  F  K  K  N  Y  H  M  Y  V  Y  P  V  H  W  Q  F  S  E  L  D
 436  Y  L  D  E  K  R  L  L  A  G  D  H  P  I  D  L  L  L  R  D  F  K  K  N  Y  H  M  Y  V  Y  P  V  H  W  Q  F  S  E  L  D
                                                                                   EF-hand 1

1801 caacaccctatggatagagtcttgacacattctgaacttgctcctctgcgagcatctctggtgcccatggaacactgcataacccgtttctttgaggagtgtgaccccaacaaggataag
 601  Q  H  P  M  D  R  V  L  T  H  S  E  L  A  P  L  R  A  S  L  V  P  M  E  H  C  I  T  R  F  F  E  E  C  D  P  N  K  D  K
 476  Q  H  P  M  D  R  V  L  T  H  S  E  L  A  P  L  R  A  S  L  V  P  M  E  H  C  I  T  R  F  F  E  E  C  D  P  N  K  D  K
      EF-hand 1                                                                EF-hand 2

1921 cacatcaccctgaaggagtggggccactgctttggaattaaagaagaggacatagatgaaaatctcttgttttga
 641  H  I  T  L  K  E  W  G  H  C  F  G  I  K  E  E  D  I  D  E  N  L  L  F  *   664
 516  H  I  T  L  K  E  W  G  H  C  F  G  I  K  E  E  D  I  D  E  N  L  L  F  *   539
      EF-hand 2
```

**Supplementary Fig.1 Taketomi et al.**

Human Hevin sequence. Upper: Q14515_1 (UniProtKB, SPRL1_HUMAN), Lower: Q14515-2 (UniProtKB, SPRL1_HUMAN), Yellow boxes indicate ASD-associated mutation sites. Red box indicates FS domain, Green box indicates EC domain.

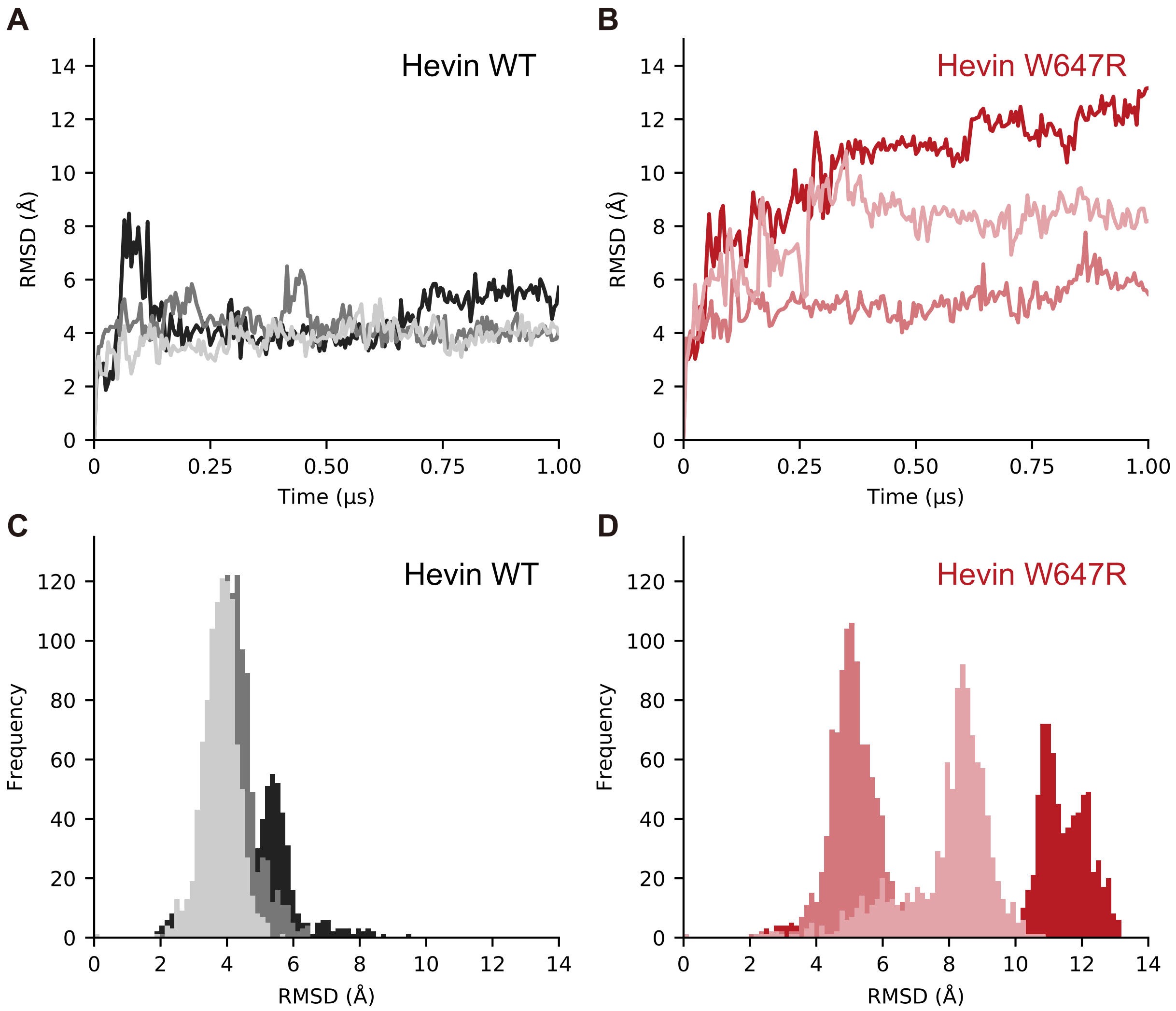


**Supplementary Fig. 2 Taketomi et al.**

RMSD profiles of wild type and W647R mutant. Time series of root-mean-square deviation (RMSD) with respect to each initial structure for 1 μs of (A) wild type and (B) W647R mutant. (C)(D) RMSD distributions of (A) wild type and (B) W647R mutant. Each color represents individual three 1-μs MD simulations.
